# Recurrent pattern completion drives the neocortical representation of sensory inference

**DOI:** 10.1101/2023.06.05.543698

**Authors:** Hyeyoung Shin, Mora B. Ogando, Lamiae Abdeladim, Severine Durand, Hannah Belski, Hannah Cabasco, Henry Loefler, Ahad Bawany, Ben Hardcastle, Josh Wilkes, Katrina Nguyen, Lucas Suarez, Tye Johnson, Warren Han, Ben Ouellette, Conor Grasso, Jackie Swapp, Vivian Ha, Ahrial Young, Shiella Caldejon, Ali Williford, Peter Groblewski, Shawn Olsen, Carly Kiselycznyk, Jerome Lecoq, Hillel Adesnik

## Abstract

When sensory information is incomplete or ambiguous, the brain relies on prior expectations to infer perceptual objects. Despite the centrality of this process to perception, the neural mechanism of sensory inference is not known. Illusory contours (ICs) are key tools to study sensory inference because they contain edges or objects that are implied only by their spatial context. Using cellular resolution, mesoscale two-photon calcium imaging and multi-Neuropixels recordings in the mouse visual cortex, we identified a sparse subset of neurons in the primary visual cortex (V1) and higher visual areas that respond emergently to ICs. We found that these highly selective ‘IC-encoders’ mediate the neural representation of IC inference. Strikingly, selective activation of these neurons using two-photon holographic optogenetics was sufficient to recreate IC representation in the rest of the V1 network, in the absence of any visual stimulus. This outlines a model in which primary sensory cortex facilitates sensory inference by selectively strengthening input patterns that match prior expectations through local, recurrent circuitry. Our data thus suggest a clear computational purpose for recurrence in the generation of holistic percepts under sensory ambiguity. More generally, selective reinforcement of top-down predictions by pattern-completing recurrent circuits in lower sensory cortices may constitute a key step in sensory inference.

## Main

Illusions arise from rational mistakes in sensory inference. For example, a rational perceptual interpretation of the Kanizsa illusion is that of a white triangle occluding three black circles (Fig. 1a)^1^. The nervous system likely evolved to rely on inference because we must frequently infer objects from partial information, such as during occlusion. Many species, including humans, non-human primates, mice, fish and even insects, perceive illusory contours (ICs), implying that sensory inference is fundamental to perception^2–7^. For example, mice trained to perform an orientation discrimination task on real edges can seamlessly generalize their performance to ICs, and vice versa^5, 6^.

**Figure 1.**
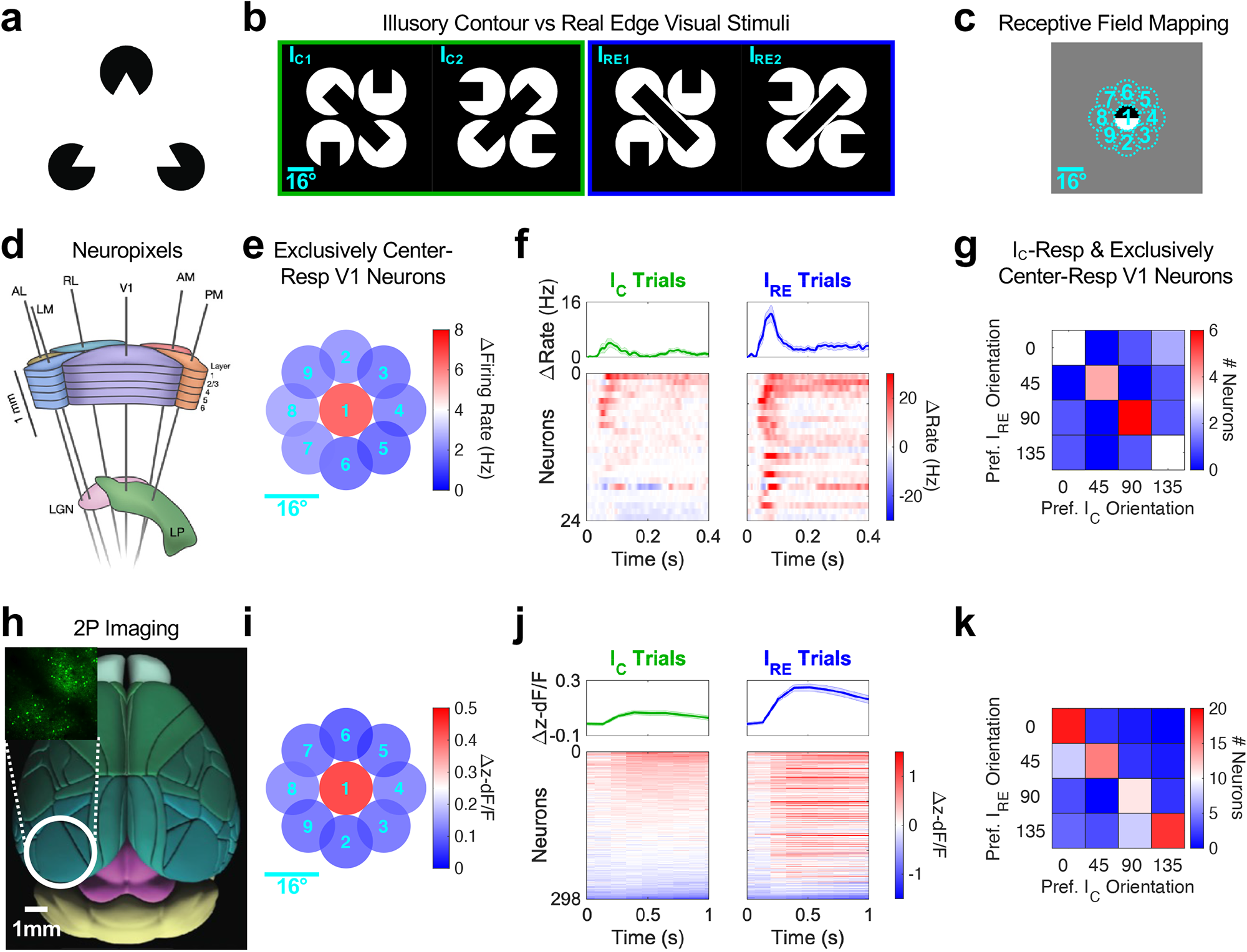
Mouse V1 neurons respond to illusory contours despite the lack of visual information within their receptive fields. **a,** Kanizsa triangle. **b,** A subset of visual stimuli used in this experiment. **c,** RF positions of each neuron was mapped using 16 degree circular grating patches, appearing in one of the 9 positions depicted. Circular patch in position 1 corresponds to the gap region between four white circles in b. **d,** Schematic of 6 Neuropixels probes insertion into V1, LM, RL, AL, PM and AM. **e,** Evoked activity at each position of V1 neurons that exclusively responded to grating patches in position 1, corresponding to the illusory gap region, and not to any of the other positions shown in c (n=24 exclusively center-responsive neurons out of 2,395 V1 neurons; 14 sessions from 14 mice). **f,** Peri-stimulus time histogram (PSTH) of exclusively center-responsive V1 neurons on I_C_ and I_RE_ trials (top: averaged, bottom: individual units). **g,** For the subset of units in f that significantly respond to I_C_ images, preferred orientation was compared between ICs (I_C_ stimuli) and real edges (I_RE_ stimuli). **h,** Schematic of the brain with areas demarcated. Inset: example 2p image of GCaMP6 expressing neurons in V1. **i-k,** Same as e-g, but for 2p imaging dataset (n=298 exclusively center-responsive neurons out of 18,576 V1 layer 2/3 neurons; 29 sessions from 5 mice).

Illusions are uniquely suited for dissociating faithful from inference-based sensory representations in the brain. Pioneering work in the primate visual cortex identified neurons that respond to ICs as if they contained real edges, despite the lack of any actual contrast in their receptive fields (RFs)^8–10^. Such responses may represent a neural correlate of emergent, or Gestalt, perception, in which the perceived whole is greater than the sum of its parts (or sensory inputs). While lower visual cortical areas are primarily faithful to image segments, higher visual areas show stronger emergent representation of the ICs^11–13^. Further, recent work in mice showed that optogenetic silencing of a higher visual area (LM) during IC presentation reduced IC-evoked responses in V1^6^. These data strongly support a model where circuits in higher cortical areas first compute the presence of ICs and then feed back these representations as predictive inferences to lower cortical areas. However, it remains unclear why these sensory predictions are fed back to V1. One possibility is that recurrent circuits in V1 are primed for neural pattern completion and thereby selectively strengthen IC inference signals arriving from higher cortical areas. In such a view, selective reinforcement of top-down predictions by pattern-completing recurrent circuits in lower cortical areas could represent a key step in Gestalt perception.

To address these questions, we used mesoscale two-photon (2p) calcium imaging, multi-Neuropixels extracellular electrophysiology recordings, and 2p holographic optogenetics in awake mice viewing images that contain ICs. Simultaneous recording from large neural populations was essential for two reasons: first, IC encoding neurons represented only a very sparse fraction of all visually responsive neurons; and second, experimentally assessing the presence of IC representations required multivariate analysis of a large unbiased population of neurons. Briefly, we asked how neural decoders trained to discriminate between IC stimuli generalized their performance to the discrimination of real edge stimuli.

Using these approaches, we found a representation of ICs in layer 2/3 of V1 and LM, but not in layer 4 of V1. The visually driven activity of only a small subset of neurons that responded emergently to the IC (‘IC-encoders’) was necessary for sensory inference, as assessed by neural decoding analysis. To test whether these IC-encoders could drive IC representations on their own, we holographically photo-activated them in the absence of any visual stimulus. We found that photo-activating V1 IC-encoders recreated IC representations via neural pattern completion. These data provide critical insight into the neocortical mechanism of sensory inference, highlighting recurrent connectivity in primary sensory areas as key to this fundamental computation of the neocortex.

### Illusory contour responses of mouse V1 neurons

To gain a comprehensive understanding for how the mouse visual cortex encodes the sensory inference of ICs, we took advantage of two high-throughput neural recording techniques with complementary strengths: Neuropixels and large-scale 2p imaging. Using these two techniques, we recorded the neural activity of awake, head-fixed mice while presenting the animals with various images containing real edges or ICs (Fig. 1b; I_C_ denotes ‘I-shaped combination’, I_RE_ denotes ‘I-shaped real edge’).

First, we asked if we could identify visual cortical neurons that respond to ICs even in the absence of any contrast within their RFs, as previously shown in primates. We carefully mapped the RFs of all recorded visually responsive neurons (or single units) from the Neuropixels recordings, and selected only those cells that responded exclusively to the central 16° region of the stimulus monitor (Fig 1c-e). Of these spatially selective cells, the great majority nonetheless responded to at least one orientation of an IC stimulus in which the illusory (or ‘gap’) region encompassed the entire RF of each neuron (Fig. 1f; out of exclusively center-responsive neurons, 79.2% neurons were I_C_-responsive, and 95.8% neurons were I_RE_-responsive). Thus, despite having no contrast within their RFs, most of these neurons responded as if the RF contained a stimulus. Importantly, there was a strong degree of overlap in the orientation preference of these neurons for real edges and ICs (Fig. 1g). This demonstrates that mouse V1 contains neurons that respond emergently to ICs in a feature selective manner. Because we used a very strict inclusion criterion for this analysis, we can confidently exclude any possibility that these cells were driven by small parts of their RF that overlap with the inducer segments of the IC stimuli. To confirm these findings with an alternative approach, we applied the same analysis to V1 neurons recorded by 2p calcium imaging (Fig. 1h). Indeed, as for electrophysiology, we could identify V1 cortical neurons that responded to IC stimuli despite no contrast within the cells’ RFs (Fig. 1i,j; out of exclusively center-responsive neurons, 32.6% neurons were I_C_-responsive, and 61.4% neurons were I_RE_-responsive). Again, for most neurons, the orientation preference for real edges matched that of ICs (Fig. 1k).

Although the classical definition of IC encoding neurons are neurons whose RF is contained entirely within the gap region of the IC^8, 9^, neurons with RFs that overlap one of the inducer segments could still detect the presence of the IC. Recent work in the mouse visual system has demonstrated that V1 cortical neurons are highly sensitive to the specific content of ‘extra-classical’ stimuli, i.e., contextual information outside of their classical RFs^14–16^. No study has yet addressed whether ICs might represent a particularly important class of such extra-classical stimuli. Thus, we speculated that we might find neurons that selectively encode ICs as an emergent, global feature of the image even if their RFs were not centered on the gap.

To test this notion, we needed to design a set of visual stimuli that would distinguish between cells that simply responded to low level features of the image *per se* (i.e., one of the inducer segments), versus the emergent IC inference. Thus, we devised a set of four stimuli that could disambiguate low-level from emergent responses (Fig. 2a). Two of the stimuli contained illusory bars (I_C1_, I_C2_), while the other two were constructed by combining the upper and lower halves of the first two images, such that the illusory bars were eliminated (‘L-shaped combination’, L_C1_, L_C2_). This design ensured that response selectivity for one of the four images cannot be explained by sensitivity to the component inducer segments; instead, it can only be explained by sensitivity to a global feature of the image. Using these stimuli, we could identify approximately 3% of neurons in V1 with selectivity for one of the two I_C_ images, and no significant response to either L_C_ image (Fig. 2b). Thus, we could define these neurons as ‘IC-encoders’, irrespective of their RFs. IC-encoders were tuned to the emergent orientation of the illusory bar (Fig. 2c). Interestingly, although some of these neurons had RFs that mapped to the gap region of the IC, the majority did not (Extended Data Fig. 1e). This indicates that IC-encoders can have their RFs anywhere in the image and still be selective to the emergent IC.

**Figure 2.**
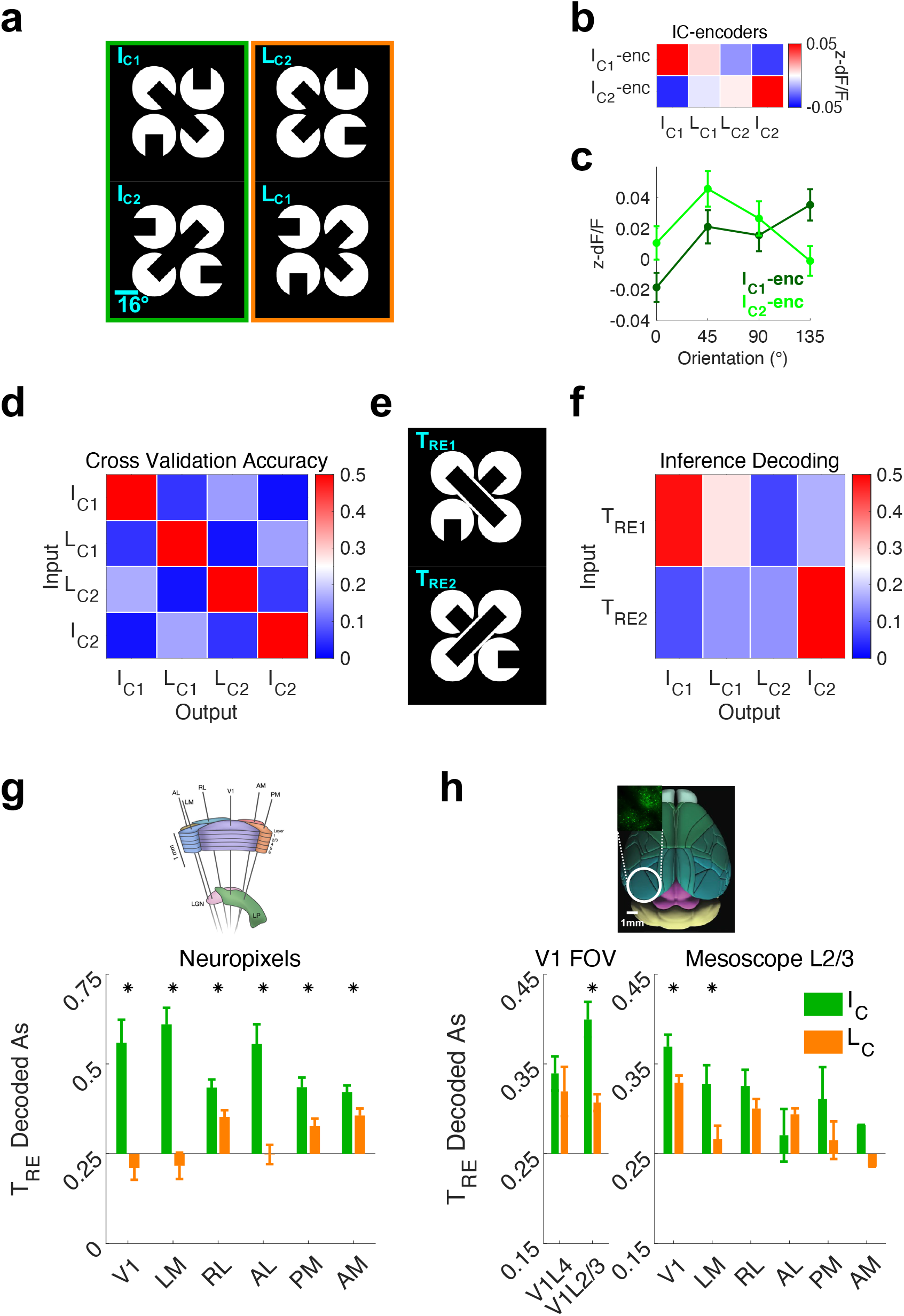
Illusory contour inference is represented in layer 2/3 of V1 and LM. **a,** Visual stimuli used for decoding analyses. L_C_ stimuli were designed such that the sum of parts for the I_C_ image pair would be equivalent to that of L_C_ image pair: L_C1_ image is constructed from bottom half of I_C1_ and top half of I_C2_, and L_C2_ is constructed as vice versa. **b,** I_C1_-encoders and I_C2_-encoders constitute IC-encoders. Figure shows average evoked responses of these neurons. **c,** Orientation tuning curves, measured with standard static gratings, averaged over each IC-encoder subgroup (mean ± SEM across neurons). **d,** 10-fold cross-validation performance of linear SVM trained on the 4 trial types shown in a. Confusion matrix shows the average decoding performance of V1 neurons in the Neuropixels dataset (n=14 sessions; average 180 V1 neurons per session, 400 repetitions per trial type). **e,** T_RE_ stimuli, which have equivalent pixel overlap with an I_C_ and an L_C_. **f,** Using the decoder described in c, neural activity evoked by T_RE_ images was classified into one of the following 4 labels; I_C1_, L_C1_, L_C2_, I_C2_ (inference decoding). **g,** Inference decoding of each visual cortical area in the Neuropixels dataset (mean ± SEM across sessions, *p<0.05, right-tailed Wilcoxon signed-rank test; n=14 sessions). **h,** Inference decoding of the 2p dataset. The proportion of T_RE_ trials that are decoded as the corresponding I_C_ (green) and L_C_ (orange) images were averaged across the two image sets (mean ± SEM across sessions, *p<0.05, right-tailed Wilcoxon signed-rank test; n=8/20/19 sessions for V1L4 FOV, V1L2/3 FOV, and mesoscope, respectively. For each visual area, sessions with less than 100 neurons in that area was discarded).

### Representation of illusory contour inference across the mouse visual cortical hierarchy

Top-down corticocortical feedback generates IC encoding in V1 in both primates and mice^6, 13, 17, 18^. To understand how ICs are encoded more broadly across the mouse visual system, we recorded from six visual areas simultaneously (V1, LM, AL, RL, AM and PM), either by targeting individual Neuropixels probes^19, 20^ to each area, or by using a 2p mesoscope^21^. We also compared V1 layer 2/3 (V1 L2/3) and layer 4 (V1 L4) using multi-plane imaging in a standard 2p microscope.

Using the same approach as above for V1, we could identify sparse populations of IC-encoders throughout these visual areas both in the Neuropixels and in the 2p mesoscope datasets (Extended Data Fig. 2a-b). However, if we simply averaged the response of all neurons in each area, we observed that images containing illusory bars (I_C_’s) did not generate more activity than those that did not (L_C_’s), in any of the six visual areas we looked at (Extended Data Fig. 2c-d). This implies that, on average, IC stimuli do not generally evoke greater activity in visual areas than non-IC stimuli.

The lack of a mean difference in the cortical response to IC stimuli, however, does not necessarily imply the lack of emergent IC encoding in the pattern of neural activity. Prior studies of IC representations in the cortex used low channel count electrodes, and thus could only record one or a small number of neurons simultaneously. These studies were thus limited to neuron-by-neuron (univariate) analyses which would miss representations that only emerge in neural activity patterns (multivariate). Owing to the high-throughput nature of our recording techniques, we could apply powerful multivariate approaches that can test more generally for IC representations. We thus trained machine learning algorithms (linear support vector machines, SVM) to classify visual evoked neural activity patterns into corresponding visual labels for the four trial types described above (I_C1_, L_C1_, L_C2_, and I_C2_ trials; chance performance is 0.25). The 10-fold cross-validation performance was significantly above chance, indicating that neural activity patterns evoked by these four images were highly distinct at the multivariate level (Fig. 2d).

To test whether IC inference is represented in the visually evoked neural activity patterns, we needed to design a new pair of images that could test whether the machine learning decoder was sensitive to the emergent ICs or was simply distinguishing the four stimuli based on lower-level image differences. To this end, we designed two new stimuli, called ‘T-shaped real edges’ (T_RE_; Fig. 2e): T_RE1_ has equal pixel overlap with I_C1_ and L_C1_ (and, correspondingly, T_RE2_ has equal pixel overlap with I_C2_ and L_C2_). Importantly, T_RE_ stimuli contain explicit real edges, unlike any of the four stimuli used to train the decoder. Thus, if visually evoked activity patterns are faithful to the images, the decoder should be equally likely to classify T_RE_-evoked neural activity patterns as I_C_ or L_C_, because they have identical pixel overlap with the T_RE_ stimulus. On the contrary, if a cortical area contains the neural representation of the IC inference, T_RE_-response in that area should be more similar to I_C_-response than to L_C_-response. To quantify this effect, we computed the difference in classification of T_RE_ as I_C_ versus L_C_ (averaging over the two image sets) and termed this analysis “inference decoding”. Applying this analysis to our Neuropixels V1 recordings showed that T_RE_ was significantly more likely to be decoded as I_C_ than as L_C_ (Fig. 2f). This result indicates the presence of the IC inference representation in mouse V1.

To survey IC inference representations across the mouse visual cortex, we applied the same analysis to each visual area. In the Neuropixels dataset, every visual area showed significant inference performance, with V1, LM and AL showing the highest performance (Fig. 2g). Similarly, L2/3 of V1 and LM showed the greatest inference performance in the 2p mesoscope dataset (Fig. 2h). Interestingly, V1 L4 did not show significant inference performance, consistent with the notion that L4 is more faithful than L2/3 to low level image features (Fig. 2h). Together, these results are consistent with a model in which the neocortical representation of ICs arises via corticocortical feedforward and feedback pathways between V1 and higher visual areas, specifically area LM. However, they shed no light on what role this feedback has, or more generally, whether V1 contributes to IC inference beyond simply reflecting the inference computed in higher visual areas.

### Neural pattern completion of illusory contour representation in V1

So far, our data has established mechanisms of IC encoding at two levels: at the univariate level, we identified the existence of a sparse population of IC-encoders, and at the multivariate level, we found the existence of neural representation of IC inference in patterns of visual cortical activity. To connect these two levels, we sought to determine if IC-encoders are necessary for the multivariate representation of IC inference, and whether they might be sufficient to recreate it through recurrent circuitry. For this we took advantage of 2p calcium imaging rather than electrophysiology since it affords the necessary dense sampling of V1 L2/3 activity. Moreover, only 2p imaging allows us to target IC-encoder ensembles for selective activation in 2p holographic optogenetics experiments.

First, to test whether IC-encoders are required for the representation of IC inference, we analyzed the change in decoder performance when zeroing out their activity in the decoder input. Indeed, this abolished IC inference performance, as measured by the decoder’s bias to classify T_RE_ images as I_C_ versus L_C_ stimuli (Fig. 3b). This demonstrates that the activity of IC-encoders is required for the neural representation of IC inference. Next, we took the converse approach and asked if the information encoded in IC-encoders alone might be sufficient to represent IC inference. To this end, we trained a new decoder only on IC-encoders (average 31/1968 neurons per session). Strikingly, despite training this decoder on the activity of dramatically fewer cells, it showed high levels of IC inference performance (Fig. 3c).

**Figure 3.**
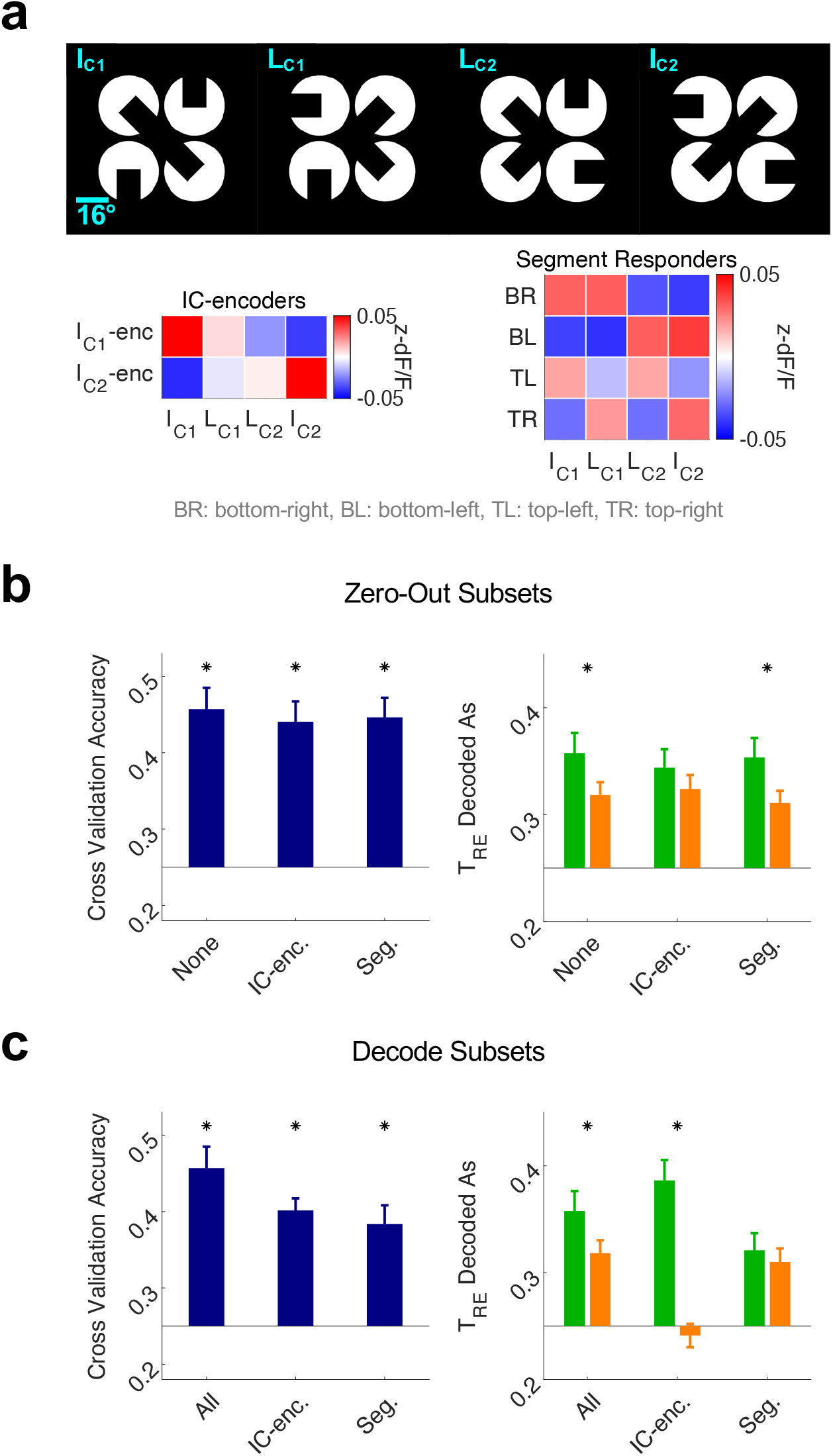
IC-encoders mediate the representation of illusory contour inference in V1 layer 2/3. **a,** Top: I_C_ and L_C_ stimuli. Bottom left: average response of neurons defined as ‘IC-encoders’ to the I_C_ and L_C_ stimuli (1.6% of ROIs in V1 L2/3 FOV, n=24 sessions from 4 mice). Bottom right: average response of neurons defined as segment responders (3.5% of ROIs). **b,** Decoder performance when zeroing out subsets of neurons, IC-encoders or segment responders, in the input to the decoder (mean ± SEM across sessions, *p<0.05, right-tailed Wilcoxon signed-rank test). **c,** Decoder performance when decoding only off of subsets of neurons (same subsets as b; mean ± SEM across sessions, *p<0.05, right-tailed Wilcoxon signed-rank test).

To test whether this result could simply be explained by the visual responsiveness of IC-encoders, we executed an analogous set of analyses on ‘segment responders’, i.e., neurons that respond to segments of the I_C_ and L_C_ images and thus do not discriminate between I_C_ and L_C_ (average 69/1968 neurons per session). Segment responders respond either to the bottom-right (BR) segment of I_C1_ and L_C1_, or the bottom-left (BL) segment of I_C2_ and L_C2_, or the top-left (TL) segment of I_C1_ and L_C2_, or the top-right (TR) segment of I_C2_ and L_C1_ (Fig. 3a, Extended Data Fig. 1b). Cross validation performance after zeroing out segment responders was not significantly different from zeroing out IC-encoders (Fig. 3b *left*), and cross validation performance when decoding subsets was also not different between segment responders and IC-encoders (Fig. 3c *left*). These results indicate that segment responders contain as much visual information as IC-encoders. Even so, zeroing out these segment responders’ activity in the original decoder preserved the IC inference performance (Fig. 3b *right*), and training a new decoder only on their activity did not yield a significant IC inference performance (Fig. 3c *right*). These results demonstrate that IC-encoders, at least at the level of post-hoc decoding analysis, are a special class of cortical neurons that are necessary for the neural representation of IC inference. Next, we sought to address whether these neurons have a direct causal role in shaping IC representation.

Prior work has shown that IC encoding in V1 arises via top-down feedback; this feedback likely depends on integration of convergent feedforward input of V1 neurons that are themselves driven in a feedforward manner by the IC inducing segments^6, 17, 18^. While this model explains how cortico-cortical circuits between V1 and higher visual areas can generate IC encoding in V1, it offers no explanation for why IC encoding should exist in V1 at all, or how it might causally contribute to IC representations.

We hypothesized that V1 IC-encoders might selectively strengthen IC representations through recurrent circuitry. Such a mechanism could promote sensory inference by preferentially fortifying input patterns that match prior expectations. To test for such recurrent pattern completion, we employed 2p holographic optogenetics to selectively photo-activate ensembles of IC-encoders. We developed an “all-optical read-write” experimental pipeline consisting of three phases (Fig. 4a). In phase one (the “read” stage), we recorded visual responses of several thousand V1 L2/3 neurons to the IC image set. In phase two (the “online analysis” stage), we analyzed the visual response properties of every neuron in the field-of-view (FOV) to identify IC-encoders. In phase three (the “write” stage), we holographically stimulated these IC-encoders (Fig. 4b), while simultaneously imaging the same FOV in the absence of visual inputs. With this approach, we could test whether selectively photo-activating ensembles of IC-encoders would be sufficient to generate the neural representation of IC inference on their own. If so, this would imply that during normal visual activity, IC-encoders reinforce IC inference.

**Figure 4.**
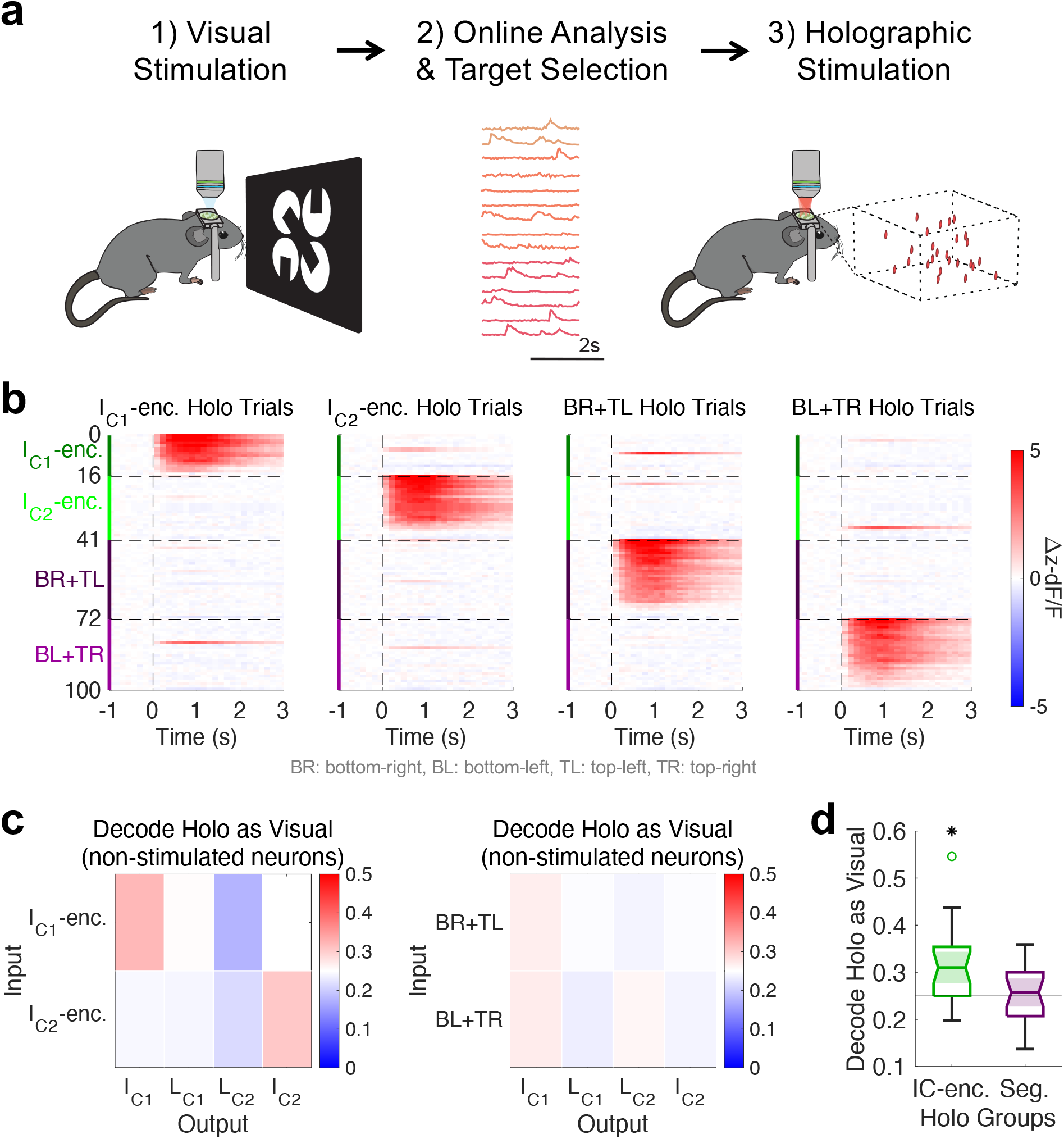
Two-photon holographic optogenetic stimulation of IC-encoders is sufficient for recurrent pattern completion of illusory contour representation in V1 layer 2/3. **a,** Experimental pipeline, consisting of three stages: First, visual responses are imaged; second, visual response properties are analyzed; third, functional subsets of neurons are stimulated via 2p holographic optogenetics (n=24 sessions from 4 mice). **b,** Holography evoked activity of targeted neurons on distinct holography trials in stage 3. Four distinct subsets of neurons are targeted; I_C1_-encoders, I_C2_-encoders, BR- and TL-segment responders, and BL-and TR-segment responders. **c,** Decoder trained on visual trials was used to classify holography evoked activity in non-stimulated neurons (neurons >50μm from all holography targets). **d,** Replotting of data in c, with significance test against chance performance of 0.25. (*p<0.05 right-tailed Wilcoxon signed-rank test across n=24 sessions).

We trained a neural decoder to identify the visual stimuli (I_C1_, L_C1_, L_C2_, I_C2_) from the visually evoked neural activity patterns, as in Fig. 2-3. We excluded any cell that could have been directly activated by the photo-stimulation (non-stimulated neurons, >50μm from all holographic targets). Non-stimulated neurons showed above-chance cross-validation accuracy.

Next, to test whether photo-activating IC encoding ensembles could recreate key aspects of IC-evoked activity patterns, we asked how the decoder, trained only on visually evoked activity, would classify non-visual holography-evoked network activity patterns. Indeed, the decoder classified IC-encoder stimulation trials as the corresponding I_C_ image with significantly above-chance probability (Fig. 4c *left*; average 18 IC-encoders were targeted). Thus, stimulating IC-encoders even in the absence of any visual stimulus was sufficient to recreate neural activity patterns that significantly resembled those visually evoked by the corresponding I_C_ image. This constitutes a form of neural pattern completion. Conversely, photo-activating a similar number of segment responders did not evoke decodable patterns (Fig. 4c *right*; average 22 segment responders were targeted). Consistent with decoding results, holographic stimulation of IC-encoders, but not segment responders, drove selective activity in I_C_-responsive neurons (Extended Data Fig. 4c). Together, these results confirm that activating IC-encoders *per se*, and not simply adding activity to visually responsive neurons, is required to generate meaningful representations in the rest of the network (Fig. 4d).

## Discussion

Here we combined large-scale electrophysiology, 2p imaging, and 2p optogenetics to establish a direct, causal link between neural pattern completion and sensory inference. First, our high-throughput techniques allowed us to identify sparse subpopulations of cortical neurons in an unbiased manner that specifically encode ICs as emergent features of the image. We discovered that these IC-encoders could have RFs both within and beyond the illusory gap region. Next, we leveraged the large numbers of simultaneously recorded neurons to develop a new multivariate approach to identify patterns of neural activity that encode sensory inferences. Using this novel ‘inference decoding’ paradigm, we found particularly strong representations of IC inference in V1 and LM across both the Neuropixels dataset and the imaging datasets in L2/3. In contrast, thalamo-recipient L4 in V1 had negligible representation of IC inference. Further, IC-encoders in V1 were required for the representation of sensory inferences. Finally, and perhaps most strikingly, we found that 2p holographic stimulation of IC-encoders could recreate the visually evoked pattern of activity in the rest of the V1 L2/3 network in the total absence of any visual stimulus. These data outline a model where reverberating activity between L2/3 of V1 and LM compute and selectively reinforce representations of inferred contours.

Our results identify V1 recurrent activity to be a critical missing step in existing models of IC encoding (Extended Data Fig. 4a). Prior literature emphasized the role of top-down feedback in generating IC representation in V1^6, 13^, positing a three-step mechanism for IC encoding^17, 18^: First, the inducer segments evoke bottom-up feedforward activity in V1 segment responders. Second, neurons in higher visual areas spatially integrate feedforward input from the V1 segment responders to generate an emergent representation of the IC. Third, top-down feedback drives V1 IC-encoders, giving rise to the representation of IC inference in V1. Although this prior model explains why IC responses emerge in V1, they don’t explain what role they might play, if any. Our findings fill in this gap by highlighting a fourth critical step within V1 L2/3: recurrent connectivity from IC-encoders to other IC responsive neurons strengthen IC representations through neural pattern completion. Finally, this recurrent activity within V1 may help ensure that only visual signals that match prior expectations are selectively reinforced during reverberating activity between V1 and higher visual areas. This reverberating activity may then propagate further downstream, eventually igniting perception^22^.

Notably, prior models of IC encoding do not explicitly posit the presence of recurrent connectivity at the neuronal level, since the neurons sending bottom-up feedforward projections from V1 to higher visual areas (segment responders) are assumed to be distinct from the neurons receiving top-down feedback projections (IC-encoders)^17, 18^. As such, our results confirm a key prediction from computational studies of artificial neural networks; that recurrent connectivity is necessary for IC inference^23, 24^. Conventional convolutional neural networks, which have purely feedforward architectures, failed to “perceive” ICs^23^, but adding in recurrent connectivity led to the emergence of IC representations^24^. Our results thus provide the first experimental evidence that recurrent connectivity within V1 has a critical role in driving IC representations.

In contexts of visual occlusion, top-down predictions about the occluded objects are also thought to be implemented by feedback connections from higher sensory areas^15, 25, 26^. Recurrent reinforcement of top-down predictions in lower sensory areas likely plays a critical role in visual object recognition during occlusion as well^27^. Thus, recurrent pattern completion may generally be an essential step in sensory inference.

Several prior studies have shown that cortical stimulation of a small number of neurons could trigger highly structured patterns of neural activity, or even goal-directed behavior, in the absence of any sensory inputs^28–31^. Such structured patterns can even emerge during spontaneous activity^32, 33^. These studies revealed that cortical circuits are primed for pattern completion, likely due to structured connectivity^34–37^. However, none of these studies observed pattern completion in the context of sensory inference. Given the sparsity of IC-encoders, it seems highly unlikely that any of these prior cortical stimulation studies would have activated them systematically. Instead, there are likely to be multiple routes to drive pattern completion in the cortex.

We propose that the neural pattern completion that underlies IC encoding is ubiquitous in active perceptual processing of the external world. We are constantly faced with incomplete or ambiguous sensory information. The mechanisms we identified here may be central for enabling object recognition despite how objects are frequently ambiguous in the retinal image.

Our work fills in a critical missing gap in cortical models of sensory inference. Our data suggests that feedback-driven recurrent activity in lower cortical areas triggers neocortical pattern completion. This recurrent pattern completion selectively reinforces top-down predictions that match prior expectations, leading to the emergence of holistic percepts despite sensory ambiguity.

## Methods

### Animals

All experiments were performed in C57/B6 mice of both sexes, aged 6 weeks and older. Imaging experiments in Figures 1-2 were conducted with CaMKII-tTA;tetO-GcaMP6s mice (V1 L2/3 and Mesoscope L2/3 data), and Scnn1a-Tg3-Cre;Ai162(TIT2L-GC6s-ICL-tTA2)-D mice (V1 L4 data). Neuropixels experiments in Figs. 1-2 were conducted with Sst-IRES-Cre;Ai32(RCL-ChR2(H134R)_EYFP) mice (n=10) and PV-IRES-Cre;Ai32(RCL-ChR2(H134R)_EYFP) mice (n=4). 2p holographic optogenetics experiments in Fig. 3-4 were conducted with AAV8-hSyn-GCaMP6m-p2A-ChRmine-Kv2.1-WPRE injected in PV-IRES-Cre;RCL-tdTomato mice (n=2) and AAV9-2YF-hSyn-DIO-GCaMP6m-P2A-ChRmine-Kv2.1-WPRE in Emx1-Cre mice (n=2). All experiments on animals were conducted with approval of the Animal Care and Use Committee of the University of California, Berkeley (imaging and 2p holographic optogenetics experiments) and Allen Institute’s Institutional Animal Care and Use Committee (Neuropixels experiments).

### Surgery

Mice were anesthetized with isoflurane (2%) and mounted on a stereotaxic apparatus. Buprenorphine (0.05mg/kg, analgesic) and dexamethasone (2mg/kg, anti-inflammatory) were injected subcutaneously. Body temperature was maintained at 36.5°C. After hair removal and disinfection with 70% ethanol and 5% iodine, scalp was removed and facia was retracted. A circular craniotomy of 3-5mm was made on the left hemisphere using a biopsy punch (Robbins Instruments) and/or a dental drill (Foredom) with a 0.5mm drill bit (FST Item No. 19007-05). Glass coverslips of 3-5mm diameter was placed on top of the craniotomy and sealed with dental cement (Metabond C&B). These glass coverslips, or ‘windows’, were made beforehand by gluing 2 layers of round cover glass (e.g., 1X 3mm + 1X 5mm, Warner Instruments, #1 thickness) with Norland Optical Adhesive #71 (UV-cured). Finally, a titanium headplate was fixed to the skull with dental cement (Metabond C&B). The dental cement was coated in a layer of black oxide to mitigate light leakage during 2p imaging.

For 2p holographic optogenetics experiments, we injected the GCaMP6m-p2A-ChRmine-Kv2.1 virus during the window implant surgery (500nL per 2 depths and 2 sites). Injection speed was 100-250 nL/min, and after injection in each site, pipette was held in place for 5 minutes. Injection sites were centered at 2.7mm lateral, 0.2mm anterior relative to lambda, and injection depths were 120µm and 250µm.

### Neuropixels Experiments

The Neuropixels data was acquired at the Allen Institute as part of the OpenScope project that allows the community to apply for observation on the Allen Brain Observatory platform (https://alleninstitute.org/division/mindscope/openscope/). The Neuropixels pipeline at the Allen Institute is described in detail in Siegle et al.^20^ and Durand et al.^38^, as well as the online white paper (https://brainmapportal-live-4cc80a57cd6e400d854-f7fdcae.divio-media.net/filer_public/80/75/8075a100-ca64-429a-b39a-569121b612b2/neuropixels_visual_coding_-_white_paper_v10.pdf). Briefly, the mouse underwent headplate implantation and craniotomy surgery, in a procedure similar to the one described above. The 5mm craniotomy was sealed with a glass coverslip, the bottom of which was coated with silicone to reduce adhesion to the brain surface. Glass coverslip held in place with Vetbond and Kwik-Cast (World Precision Instruments) until the day of the recording (several weeks later). Intrinsic signal imaging was performed to identify visual area boundaries^39^. Based on the identified visual areas, an insertion window was designed for each mouse. Mice were habituated to head-fixation and visual stimulation over a period of two weeks.

On the day of recording, the cranial coverslip was removed and replaced with the insertion window containing holes aligned to six cortical visual areas. Mice were allowed to recover for 1-2 hours after insertion window placement, before being head-fixed in the recording rig.

Six Neuropixels probes were targeted to each of the six visual cortical areas (V1, LM, RL, AL, PM, AM). Probes were doused with CM-DiI (1 mM in ethanol; Thermo Fisher, V22888) for post hoc ex vivo probe localization. Each probe was mounted on a 3-axis micromanipulator (New Scale Technologies). The tip of each probe was aligned to its associated opening in the insertion window using a coordinate transformation obtained via a previous calibration procedure. The XY locations of the six visual area targets were translated into XYZ coordinates for each 3-axis manipulator using a custom Python script. The operator then moved each probe into place with a joystick, with the probes fully retracted along the insertion axis, approximately 2.5mm above the brain surface. The probes were then manually lowered one by one to the brain surface until spikes were visible on the electrodes closest to the tip. After the probes penetrated the brain to a depth of around 100μm, they were inserted automatically at a rate of 200μm/min (total of 3.5mm or less in the brain). After the probes reached their targets, they were allowed to settle for 5–10min.

Neuropixels data was acquired at 30 kHz (spike band, 500-Hz high-pass filter) and 2.5 kHz (LFP band, 1,000-Hz low-pass filter) using the Open Ephys GUI (Siegle et al 2017 J. Neural Eng). Videos of the eye and body were acquired at 30 Hz. The angular velocity of the running wheel was recorded at the time of each stimulus frame, at approximately 60 Hz.

The spike-band data is median-subtracted, first within-channel to center the signal around zero, then across channels to remove common-mode noise. The median-subtracted data file is sent to the Kilosort2 MATLAB package (https://github.com/mouseland/Kilosort2)^40^, which applies a 150 Hz high-pass filter, followed by whitening in blocks of 32 channels. Kilosort2 models this filtered, whitened data as a sum of spike ‘templates’. The shape and locations of each template is iteratively refined until the data can be accurately reconstructed from a set of N templates at M spike times. Finally, Kilosort2 output is curated to remove putative double-counted spikes and units with artefactual waveforms. All units not classified as noise were packaged into Neurodata Without Borders (NWB) files and analyzed in this paper.

### Two-Photon Imaging Experiments

2p imaging experiments presented in Figs. 1 – 2 was conducted in two setups. V1 L2/3 and V1 L4 imaging was done on a 2p microscope (Neurolabware) with a Ti:sapphire laser (Chameleon Ultra II, Coherent), an electrotunable lens for multi-plane imaging (Optotune), 8 kHz galvo-resonant scanner, and a Nixon 16X magnification 12.5 mm focal length 0.8 NA water immersion objective. Typical image settings were as follows: FOV size 1223 X 675 μm^2^ (796 X 512 pixels), 4 planes spaced 50 μm apart sampled at volumetric frame rate 7.715 Hz, imaging wavelength 930 nm.

Mesoscope 2p imaging was done on 2p random access mesoscope (2P-RAM, Thorlabs)^21^, equipped with a Ti:sapphire laser (Mai Tai, Spectra Physics), remote focusing unit consisting of an objective and a mirror mounted on a voice coil, 12 kHz galvo-resonant scanner and Jenoptiks/Thorlabs water immersion objective with 21 mm focal length and 0.6 excitation NA and 1.0 collection NA. Typical image settings were as follows: FOV size 3031 X 2965 μm^2^ (3040 X 1484 pixels), frame rate 3.139 Hz, imaging wavelength 920nm.

During post-hoc analysis, motion correction and source extraction were done using Suite2p^41^. Suite2p output putative regions of interest (ROIs) that correspond to neuronal somas, and these ROIs were manually inspected to keep only those that looked like cell bodies. Neuropil signal was subtracted from each ROI with a coefficient of 0.7 (F_c_ = F_ROI_ – 0.7 X F_neu_). dF/F was calculated over each continuous block of recording as (F_c_-F_0_)/F_0_, where F_0_ was the mode / median of F_c_ distribution for standard / mesoscope 2p imaging, respectively. z-dF/F was calculated by z-scoring the dF/F trace across each continuous block for each ROI. Δz-dF/F was calculated by subtracting spontaneous period activity (30s at the beginning and end of each visual block).

### Two-Photon Holographic Optogenetics Experiments

For 2p holographic optogenetics experiments presented in Fig. 3-4, 3D scanless holographic optogenetics with temporal focusing (3D-SHOT) path was implemented in a Sutter MOM (Movable Objective Microscope, Sutter Instrument Co.), as described previously^42–46^. Ti:sapphire laser (Chameleon Ultra II, Coherent) was used for two-photon imaging, and femtosecond fiber laser (Satsuma HP2, 1030 nm, 2MHz, 350 fs, Amplitude Systems) was used for two-photon holographic stimulation. The holography path included a blazed diffraction grating for temporal focusing (600l/mm, 1000nm blaze, Edmund Optics 49-570 or 33010FL01-520R Newport Corporation), and a rotating diffuser to randomize the phase pattern and expand the beam. Both the imaging path and the holography path had spatial light modulators (SLM; HSP1920, 1920 X 1152 pixels, Meadowlark Optics). The imaging path SLM enabled optical axial focusing. The SLM was conjugated to the X resonant galvo and optical axial focusing was achieved by loading Fresnel-lens-like phase patterns on the SLM for each desired axial depth. The high refresh rate provided by the overdriven SLM allowed the acquisition speed to only be limited by the number of planes (no settling time or frame drop). The holography path SLM was used to display the holographic phase mask, calculated using the Gerchberg-Saxton algorithm to place several cell-sized diffraction-limited spots in 3D target positions. A zero-order block was placed in the holography path. The imaging path and the holography path were merged by a polarizing beam splitter before the microscope tube lens and the objective (Olympus 20X magnification 9mm focal length 1.0 NA water immersion objective). To limit imaging artefacts introduced by the femtosecond laser, the femtosecond laser output was synchronized to the scan phase of the galvo-resonant scanner (8 kHz) using an Arduino Mega (Arduino), gated to be only on the edges of every line scan.

Precise alignment of the imaging path and the holography path is achieved by a calibration procedure described previously^42–44^. Briefly, we place a thinly coated fluorescent slide on a substage camera to image both the imaging planes and the holograms at parametrically varied SLM coordinates. Then, the 3D coordinate transformation from imaging coordinates on each plane to SLM coordinates is computed through the ‘polyfitn’ function in Matlab.

Before each experiment, the alignment is confirmed for each remote-focused plane by “burning holes” on a fluorescent slide (photobleaching). The intended imaging coordinates are displayed on ScanImage (Vidrio) as Integration ROIs. The match between the intended coordinates and the burnt hole positions confirms the validity of the current calibration.

As described in the Results section, the “all-optical read-write” 2p holographic optogenetic experiments were conducted in three stages. First was the ‘read’ stage, where we image visual responses of neurons in the V1 L2/3 FOV. Second was the online analysis stage, where we ran Suite2p^41^ on the imaging data collected in the first stage and identified functional groups of neurons (IC-encoders, segment responders). Third was the ‘write’ stage, where we holographically stimulated these functionally identified neuron groups. Throughout all stages of the experiment, the mouse was head-fixed under the objective, and the imaging conditions were kept constant (same FOV of size 980 X 980 μm^2^, same set of 4 planes spaced 30 μm apart, with each plane being imaged every 4 frames at 7.525Hz volumetric frame rate, imaging wavelength of 920nm).

For fast online analysis, we converted ScanImage Tiff files into four h5 files, generating one h5 file per plane. Then, we ran Suite2p for each h5 file independently, in parallel (‘multiprocessing’ package in python). On an 8-core CPU (Intel Core i7 10700K 3.8GHz) with 128GB RAM, online Suite2p of a 25 min imaging data (512 X 512 pixels) took eight minutes. Then, all ROIs identified by the Suite2p were analyzed in Matlab to identify functional groups of neurons (IC-encoders and segment responders). Finally, Suite2p output ‘stat.med’ was used as the coordinate of the ROIs of interest. These Suite2p coordinates were converted to SLM coordinates per the calibration described above.

After determining the holography groups with online analysis, holograms containing all targets in each group were calculated using the Gerchberg-Saxton algorithm. Mean number of targets in each holography group were: 17 I_C1_-encoders, 18 I_C2_-encoders, 22 BR+TL-segment responders 22 BL+TR-segment responders. The holograms were diffraction efficiency corrected to achieve homogeneous power delivery of 7mW to all targeted cells. Holographic stimulation on each trial consisted of 10 pulses at 10 Hz with 10 ms pulse width, and the inter-trial interval was 4.5s. Stimulation of each holography group was repeated 50 times.

During post-hoc offline analysis, Suite2p was run over the entire recording, including Stage 1 (read stage, visual block) and Stage 3 (write stage, holography block). On average, 1968 ROIs were classified as putative neurons in the 4-plane volumetric FOV. Targets were identified as the ROI closest to each target coordinate. The target was considered valid only if the closest ROI was within 10µm, which left 81.5% of targets (mean number of valid targets for each holography group was 15 I_C1_-encoders, 16 I_C2_-encoders, 18 BR+TL-segment responders, 17 BL+TR-segment responders). Of these, holographic stimulation evoked significant activation in 91.6% of valid targets (mean number of stimulated targets for each holography group was 14 I_C1_-encoders, 14 I_C2_-encoders, 16 BR+TL-segment responders, 16 BL+TR-segment responders).

### Visual Stimulation Monitor Placement and Retinotopy

All Neuropixels recordings were according to the standardized Neuropixels visual coding pipeline. Visual stimuli were generated using custom scripts based on PsychoPy62 and were displayed using an ASUS PA248Q LCD monitor, with 1920 X 1200 pixels (55.7 cm wide, 60 Hz refresh rate).

Stimuli were presented monocularly, and the monitor was positioned 15 cm from the right eye of the mouse and spanned 120° × 95° of visual space. Monitor placement was standardized across rigs such that mouse right eye gaze would typically be directed at the center of the monitor.

For all 2p experiments, visual stimuli were generated using custom Matlab scripts based on Psychtoolbox-3^47^. In the standard 2p imaging setup, 19 inch Samsung monitor was placed 15cm from mouse right eye (1280 X 1024 pixels, 37.6 X 30.1 cm width and height, 60 Hz refresh rate). In the mesoscope 2p imaging setup and the 2p all-optical read-write setup, Adafruit Qualia 9.7’ DisplayPort Monitor was placed 8 cm from the right eye of the mouse (2048 X 1536 pixels, 19.7 X 14.8 cm width and height). The backlight of the Adafruit monitor was triggered by the galvo-resonant scanner such that the monitor would emit light only during the mirror turnaround time, similar to the gating of the holographic stimulation described above (12 kHz in the mesoscope 2p imaging setup, 8 kHz in the 2p all-optical read-write setup). In addition, in all 2p experiments, electrical tape was applied between the objective and the mouse’s headplate to prevent monitor light from contaminating 2p imaging.

In all 2p experiments, retinotopic mapping was used to place the monitor such that the most common RF of the FOV would align with the center of the monitor. In the mesoscope 2p imaging, retinotopic mapping was additionally used to determine visual area boundaries using visual field sign maps^48, 49^. After settling on the imaging FOV at the beginning of each 2p experiment, retinotopic mapping was conducted with 16° rectangular drifting grating patches appearing in one of 25 positions tiling an 80° X 80° visual degree grid. These patches had spatial frequency of 0.04 cycles/degree and a temporal frequency of 2 Hz; cycling through 8 directions (0/45/90/135/180/225/270/315°) over the course of 2 s, followed by a 1 s gray screen inter-trial interval (ITI). Each patch position was presented with 3 – 5 repeats, with randomized trial order.

### Visual Stimulus

After retinotopic mapping, visual stimulation consisted of 3 blocks; IC block, RF mapping block, size tuning block. Trial order was randomized within each block.

In the IC block, images described as I_C_, L_C_, I_RE_, T_RE_ were shown (IC configuration 1). In Neuropixels experiments and standard 2p imaging experiments in Fig. 1, these images were also shown rotated 315° clockwise in a separate block (IC configuration 0). The diameter of the white circles was 30° (visual degrees), and the distance between the centers of the diagonally placed white circles were 46° (i.e., the gap between diagonal circles were 16°). The black bar length on each inducer segment was 16°, such that the illusory contour support ratio in I_C_ images would be 2/3. Throughout each IC block, the four white circles stayed in place even during the ITI. In Neuropixels recordings, each image was presented for 0.4 s followed by 0.4 s ITI. In IC configuration 1, each image was repeated at least 50 times, and I_C_ and L_C_ images were repeated 400 times; in IC configuration 0, each image was repeated 30 times. In standard 2p recordings in Fig. 1, each image was presented for 1 s followed by a variable ITI of 1 – 2 s. Each image was repeated 10 times in 9 sessions, and 50 times in 20 sessions. In standard and mesoscope 2p recordings in Fig. 2, each image was presented for 1 s followed by 0 s ITI. Each image was repeated at least 50 times, and I_C_ and L_C_ images were repeated 400 – 500 times. In 2p holographic optogenetics experiments, each image was presented for 1 s followed by 0 s ITI. Each image was repeated 50 – 100 times.

In the RF mapping block, circular patches of drifting gratings with a fixed diameter of 16° were presented in 9 different locations, as shown in Fig. 1c; one patch was positioned at the center of the monitor, the rest were positioned at 16° distance from the center in 6/4.5/3/1.5/12/10.5/9/7.5 o’clock positions. Drifting gratings had a spatial frequency of 0.04 cycles/degree and a temporal frequency of 2 Hz. The drifting gratings had 100% contrast and were presented on a gray background. On each trial, a circular patch of drifting grating appeared in one of 9 positions and cycled through several directions spaced 45° apart, drifting in each direction for 0.25 s (4 / 8 directions lasting 1 / 2 s in Neuropixels / 2p experiments, respectively). Presentation at each location was repeated 10 times, with 0 s ITI.

In the size tuning block, circular patches of drifting gratings were presented with a fixed position at the center of the monitor, in different sizes of 0/4/8/16/32/64°. Spatial frequency, temporal frequency and contrast of drifting gratings were identical to the RF mapping block. On each trial, a circular patch of drifting grating at a given size appeared in one of 8 directions (0/45/90/135/180/225/270/315°). In Neuropixels experiments, the presentation lasted 0.25 s followed by 0.5 s ITI, and each combination of size and direction was repeated 8 times. In 2p imaging experiments, the presentation lasted 1 s followed by 0 s ITI, and each combination of size and direction was repeated 10 times. The size tuning block was skipped in 2p holographic optogenetics experiments.

### Eye Tracking Analysis

For results presented in Fig. 1e-g, we limited the analysis to trials where the mouse was fixating (fixed-gaze trials). Video of mouse eye during the neural recording session was analyzed using DeepLabCut^50^. Resting pupil position was determined as the mode of the pupil position. Fixed-gaze trials were defined as trials where pupil position was within eight visual degrees of the resting pupil position throughout the duration of the trial.

### Definition of Functional Subsets of Neurons

Exclusively center-responsive neurons were defined as neurons that exclusively responded to circular grating patches in the center position, corresponding to the illusory gap region, and not to any of the other adjacent positions (Fig. 1; Wilcoxon rank sum test between trials with circular grating patches in each position and gray-screen trials, Bonferroni-Holm correction for multiple comparisons).

IC-encoders were defined as neurons that respond to one of the I_C_ images, and to neither of the L_C_ stimuli (Kruskal-Wallis test across responses to ‘blank’ (four white circles) vs I_C1_ vs L_C1_ vs L_C2_ vs I_C2_ with Tukey-Kramer correction for multiple comparisons).

Segment responders were defined as neurons that selectively responded to inward-pointing inducer segments in each inducer position. For example, Bottom-right (BR) segment responder is defined as neurons that respond significantly more to BR_in_ than BR_out_ (Extended Data Fig. 1; Wilcoxon rank sum test).

### Decoding Analysis

For all decoding analyses presented in main figures, we used a support vector machine (SVM) with a linear kernel. First, we constructed the trial-by-trial response matrix (#Trials X #Neurons) as the response averaged over the visual stimulus presentation period (firing rate for Neuropixels data, dF/F for 2p data). We divided the trials with the training trial types (I_C1_/L_C1_/L_C2_/I_C2_) into a training set (90%) and a testing set (10%). For 10-fold cross-validation, there were 10 splits of training/testing sets, and the 10 sets of test trials were non-overlapping. On each of the 10 iterations, the trial-by-trial response matrix was z-scored based on the training set, i.e., response matrix was subtracted by the mean of training trials and divided by the standard deviation of training trials. On each iteration, SVM was trained using the ‘fitcecoc’ function in Matlab. Cross-validation accuracy was computed as the SVM’s prediction accuracy on the testing set, averaged across 10 iterations. The inference decoding performance and the holography evoked activity decoding performance were also measured as the average prediction performance of the 10 SVM’s.

The holography evoked response matrix, used as the input to the decoder in Fig. 4c-e, was calculated as the normalized dF/F in the 1 s period following the onset of the holographic stimulation. Normalization entailed subtracting the mean and dividing by the standard deviation of 1 s prestimulus baseline period across all holography trials. To analyze the effects of photoactivation in the rest of the network, we analyzed only the non-stimulated neurons that were at least 50μm away from all holographic targets. As shown by the physiological point spread function (PPSF, Extended Data Fig. 4b), all direct influence of holographic light dissipates by 50μm.

In Extended Data Fig. 3, we compared the decoding performance of several different kinds of decoders. SVMs were trained with different kernels; linear vs quadratic polynomial vs radial basis function. In addition, a fully connected artificial neural network (ANN) was trained using Pytorch^51^, similar to the decoding described in Stringer et al.^52^ The ANN had an input layer with number of nodes matching the number of neurons, two hidden layers each with 100 nodes, and one output layer with number of nodes matching the number of training trial types (i.e., 4 nodes, each corresponding to I_C1_/L_C1_/L_C2_/I_C2_). Softmax function was applied to the output layer. The ANN was trained for 50,000 epochs with Adam optimizer and CrossEntropyLoss loss function. All decoders underwent 10-fold cross-validation, as described in detail above.

## Data and Code Availability

All code and data will be made publicly available upon publication.

## Author Contributions

H.S. and H.A. conceived of the project. H.S. performed all 2p experiments, including 2p imaging, 2p mesoscope imaging and 2p holographic optogenetic experiments. M.B.O. assisted with the design and execution of 2p holographic optogenetic experiments. L.A. assisted with 2p mesoscope imaging experiments. S.D., H.B., H.C., H.L., and B.H. performed Neuropixels recordings. A.B., B.H, and J.L. preprocessed the Neuropixels data and packaged it in the NeuroData Without Borders (NWB) format. J.W., K.N., L.S., T.J., W.H., B.O., C.G. performed surgery for Neuropixels experiments. V.H., A.Y., S.C. performed intrinsic signal imaging for Neuropixels experiments. S.C., A.W., P.G., S.O., C.K. and J.L. managed various aspects of the Neuropixels experiments. J.L. managed all aspects of the OpenScope collaboration. H.S. and H.A. wrote the paper, with input from other authors.

## Acknowledgments

We thank Doris Tsao, Josh Siegle and members of the Adesnik lab, in particular Daniel P. Mossing, Nikhil Bhatla and Uday K. Jagadisan, for comments and discussions. This work was funded by the National Institutes of Health (NIH) grants U19NS107613 and Chan Zuckerberg Biohub Investigor Award to H.A. and Weill Neurohub Fellowship to H.S.

The Neuropixels dataset was obtained at the Allen Brain Observatory as part of the OpenScope project, which is operated by the Allen Institute, Mindscope program. OpenScope was supported by the NIH grant U24NS113646. We also thank the Allen Institute founder, Paul G. Allen, for his vision, encouragement, and support.

---

**Extended Data Figure 1.**
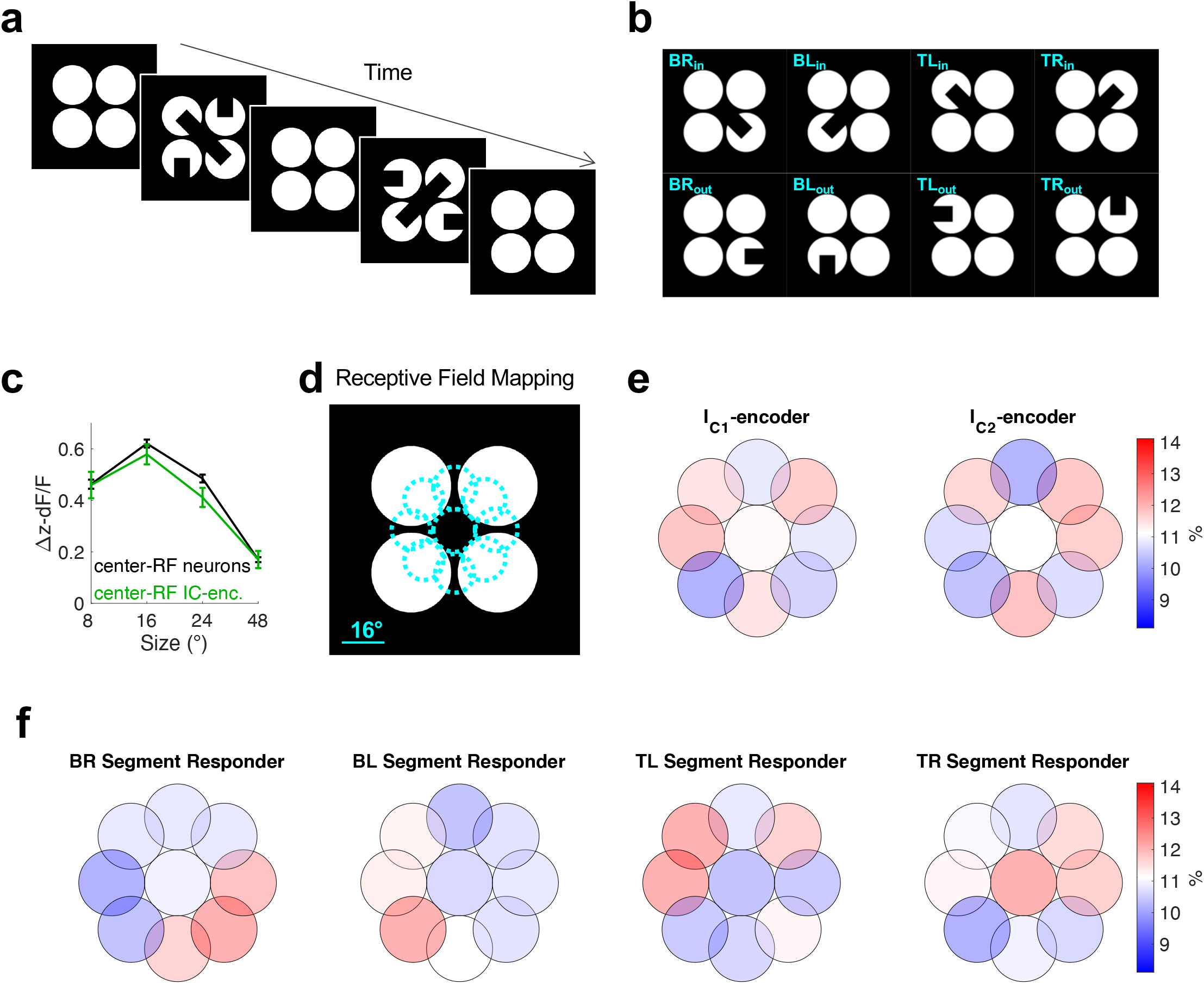
IC-encoders’ receptive fields are not biased towards the illusory gap region. **a,**Experimental schematic. Throughout each IC visual block, the four white circles stayed in place. **b,** Visual stimuli used for defining segment responders. For example, bottom-right (BR) segment responder was defined as neurons that has significantly larger responses to BR_in_ compared to BR_out_. **c,** Size tuning of V1L23 neurons (black), and IC-encoders (green) (mean ± SEM across neurons). Circular patches of drifting gratings were shown in various sizes only in the center position. Therefore, the size tuning analysis was limited to neurons with RF in the center. IC-encoders are surround suppressed to the same extent as the general population. **d,** Positions of the circular patches of drifting grating used for RF mapping (see also Fig. 1c). Center position coincides with the illusory gap region. **e,** RF position histograms of IC-encoders, showing that they are not biased towards the illusory gap region. **f,** RF position histograms of segment responders, showing that they are biased towards the image segments that are used to define them.

**Extended Data Figure 2.**
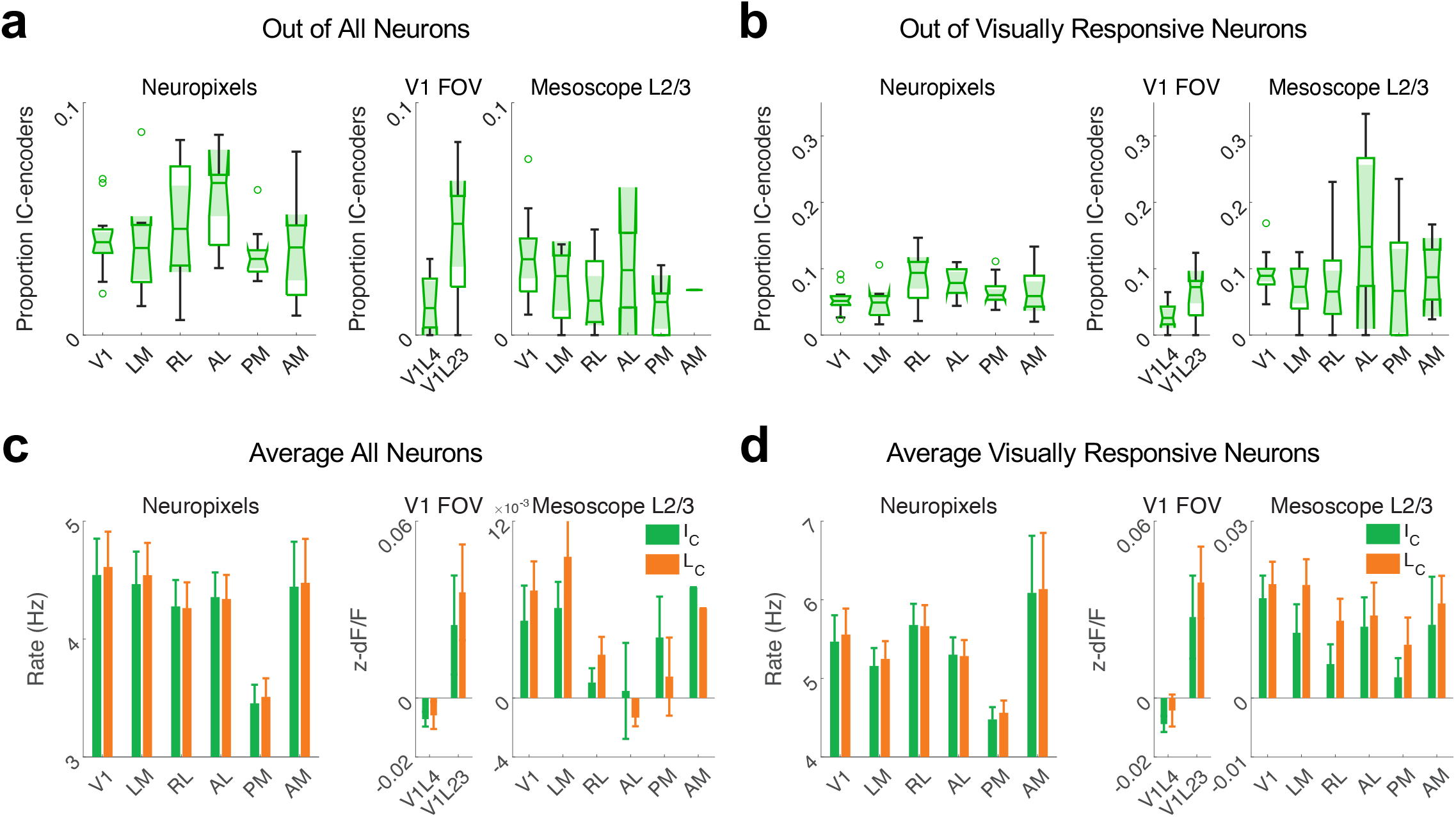
Images with illusory bars (I_C_) do not evoke greater responses than images without (L_C_). **a,** Proportion of IC-encoders, out of all neurons, in each area. **b,** Proportion of IC-encoders, out of visually responsive neurons, in each area. Visually responsive neurons are defined as neurons that respond to at least one of I_C_/L_C_ images (p<0.05 Kruskal-Wallis test across responses to ‘blank’ (four white circles) vs I_C1_ vs L_C1_ vs L_C2_ vs I_C2_). **c-d,** Response on I_C_ vs L_C_ trials, averaged across all neurons (c) or visually responsive neurons (d) for each session (mean ± SEM across sessions, *p<0.05 right-tailed Wilcoxon signed-rank test). I_C_ images do not evoke greater responses than L_C_ images.

**Extended Data Figure 3.**
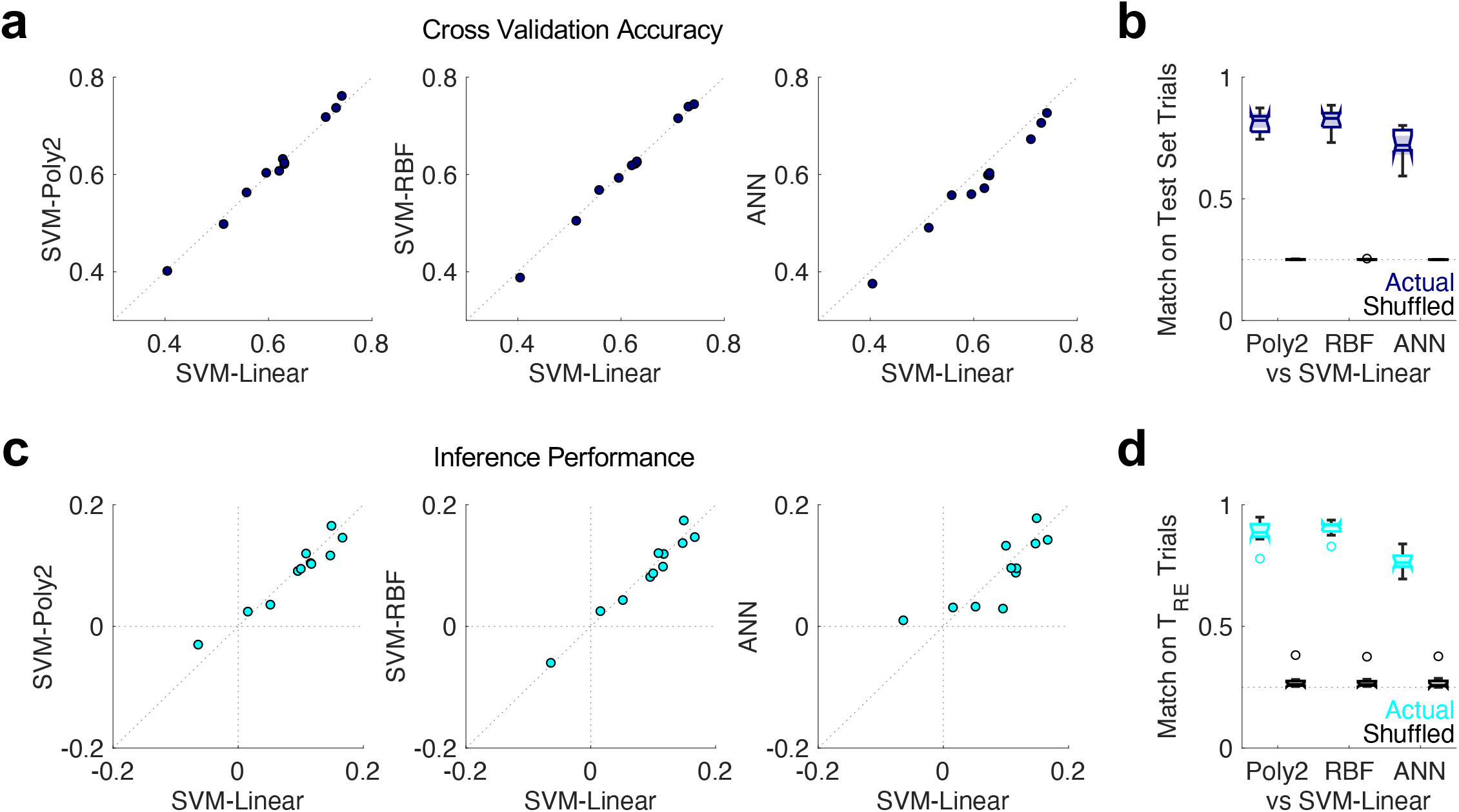
Decoder prediction is consistent across decoder types. **a,** Cross validation accuracy comparison across sessions, between SVM with a linear kernel (SVM-Linear, x-axis) vs SVM with a quadratic polynomial kernel (SVM-Poly2, y-axis on left panel), or radial basis function kernel (SVM-RBF, y-axis on center panel), or fully connected artificial neural network (ANN, y-axis on right panel). **b,** Proportion of trials in the held-out test set that have matching decoder predictions, between SVM-Linear decoder and the other three decoder types. For comparison, match proportion was also calculated with the trial order shuffled; for each session, match proportions were averaged across 1000 shuffles. **c,** Inference performance comparison across sessions, where inference performance is defined as P(T_RE_→I_C_) – P(T_RE_→L_C_). **d,** Proportion of T_RE_ trials that have matching decoder predictions.

**Extended Data Figure 4.**
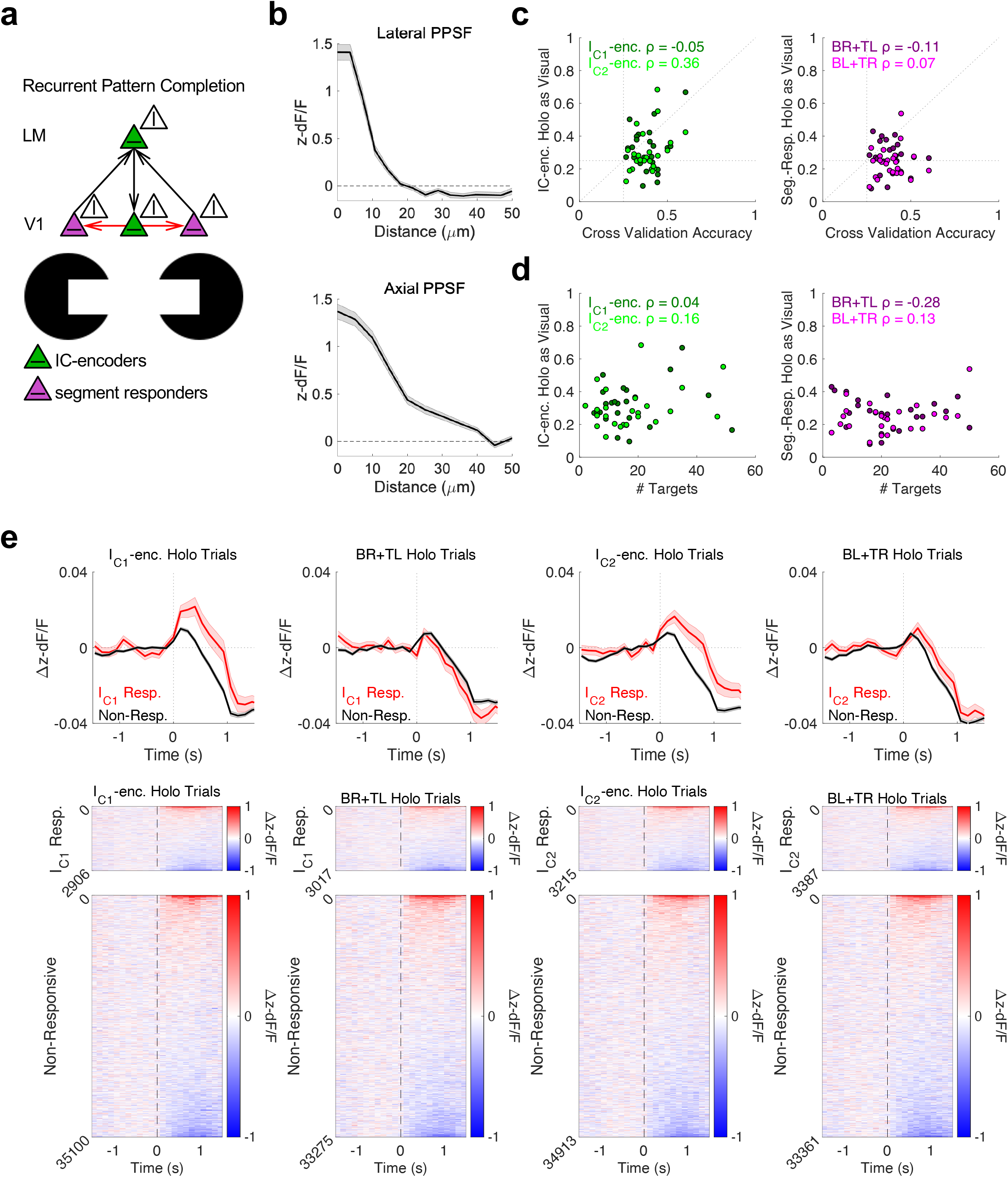
Holography evoked effects. **a,** Schematic of how IC representation arises in the visual cortical hierarchy. Black arrows show connectivity posited in prior literature^17, 18^. Red arrows are supported by our finding that photoactivation of IC-encoders recreates activity patterns visually evoked by the I_C_ images in the rest of the network (Fig. 4c-d). **b,** Physiological point spread function (PPSF) is measured by parametrically moving the holographic target position away from the center of the soma and measuring the targeted neurons’ z-dF/F. For axial PPSF, + direction indicates increasing depth. **c,** Relationship between cross-validation accuracy of decoders trained to discriminate visual trial types (I_C1_ vs L_C1_ vs L_C2_ vs I_C2_) and the proportion of holography trials decoded as corresponding visual trials. Decoder was trained on non-stimulated neurons, as described in Fig. 4. Each data point represents one session. **d,** Relationship between number of targets in each hologram and the proportion of holography trials decoded as corresponding visual trials. **e,** Holography evoked responses in non-stimulated neurons, plotted separately for neurons that are visually responsive to the I_C1_ images (red) vs non-responsive neurons (black). Both groups contain a very sparse number of driven neurons, and a larger number of suppressed neurons. On holography trials where IC-encoders were stimulated, I_C_-responsive neurons appear more driven than non-responsive neurons, consistent with the notion that IC-encoders drive neural pattern completion. While this effect is significant when pooling across neurons, the effect is not significant across sessions, suggesting that multivariate analysis with decoders is a more robust indicator of neural pattern completion.

